# Combined cellular and proteomics approach suggest differential processing of native and foreign vibrios in the sponge *Halichondria panicea*

**DOI:** 10.1101/2024.11.11.623018

**Authors:** Angela M. Marulanda-Gomez, Benjamin Mueller, Kristina Bayer, Mohammad Abukhalaf, Andreas Tholey, Sebastian Fraune, Lucia Pita, Ute Hentschel

## Abstract

Phagocytosis is a conserved cellular mechanism for food uptake, defense and general animal-microbe interactions in metazoans. How the discrimination and subsequent processing of different microbes in marine invertebrates is facilitated remains largely unknown. Thereto, we combined a recently developed phagocytic assay with proteomics analysis to compare the phagocytic activity of the sponge *Halichondria panicea* upon encounter with a native (i.e., *H. panicea* isolate; Hal 281) and a foreign (i.e., *Nematostella vectensis* isolate; NJ 1) *Vibrio*. The sponge cell fraction was recovered after *Vibrio* exposure of 30 and 60 min and used for cellular (fluorescence-activated cell sorting and microscopy) and proteomics analyses. While the number of phagocytic active cells was similar between the isolates (*p* = 0.19), the distribution of vibrios over cell types differed (*p* = 0.02) with the tendency for a higher accumulation of foreign vibrios in small phagocytic cells than the native vibrios. Initially, both vibrios elicited a mitochondrial proteomic response (e.g., TBXASI, CASP7), whereas after 60 min differentially abundant proteins were dominated by lysosomal proteins (e.g., AGA, FUCA, SCARB2) in response to the native compared to immune-related proteins (e.g., AQP9, H2A, SAMHDI) in response to the foreign vibrio. *H. panicea* may discriminate native and foreign vibrios post-cellular internalization into choanocytes, followed by differential cellular and molecular processing: Native vibrios are readily transferred to archeocytes and elicit a digestive response, whereas foreign vibrios further accumulate before being transferred and provoke an immune response. These findings provide a mechanism for immune specificity in sponges.

**Importance:** Metazoans recognize and discriminate between native (i.e., associated and/or commonly encountered) and foreign microbes. In marine invertebrates, the mechanisms of microbial discrimination and immune specificity are not well understood. Phagocytosis is a conserved cellular process from amoeba to humans that facilitates the ingestion and digestion of microbial cells and likely plays a role in this discrimination. To elucidate the molecular and cellular basis of immune specificity in marine invertebrates, we examined the differential phagocytic processing of native and foreign Vibrio isolates in a marine sponge. Our findings revealed that both vibrios provoke a generalist mitochondrial protein-mediated response, while native vibrios trigger a digestive response, and foreign vibrios elicit an immune response after being internalized into sponge cells. This study uncovers a mechanism for immune specificity in an early-divergent metazoan and provides a valuable model for studying the evolution of immunity and its role in animal-microbe interactions.

## Introduction

Phagocytosis is believed to have emerged as a means for nutrient acquisition and reallocation in unicellular amoeba. In eukaryotic organisms it was subsequently adapted in specialized immune cells, like macrophages, as a cell-defense mechanism to safeguard the organisms from potentially hazardous agents (e.g., pathogens, damaged, senescent, or apoptotic cells) and to ensure its homeostasis (1–4). Phagocytosis is a cellular process for ingesting and digesting large particles (typically > 0.5 µm in diameter), including microbes, foreign substances, and apoptotic cells, into cytosolic, membrane-bound vacuoles (i.e., phagosomes). This process comprises several subsequent steps including: Particle recognition via receptors, reorganization of the actin cytoskeleton for particle internalization, and intracellular digestion of the target particle through the activity of lytic enzymes (reviewed by (3,5)). Bacterial pathogens often targeted or manipulated phagocytosis. For example, intracellular bacterial pathogens like *Legionella pneumophila* and *Mycobacterium tuberculosis* can circumvent host phagocytosis by evading ingestion, interfering with phagosome maturation, resisting degradation, and even escaping from the phagosome (6,7). Interestingly, endosymbionts use analogous mechanisms to pathogens that aid in their colonization, acquisition, and maintenance within their host (8,9).

Many metazoans form close relationships with symbiotic microorganisms, including archaea, bacteria, fungi, and viruses (10,11). These relationships are often essential for the host organism by providing nutrients, stimulating development and growth, or protecting against pathogens (12–15). Evidence for the role of phagocytosis in microbe discrimination in endosymbiosis emerged from studies with a handful of metazoan model organisms (e.g., *Aiptasia* [anemone], *Euprymna scolopes* [squid], and *Acyrthosiphon pisum* [aphid]), for which the cultivation of symbionts, genetic manipulation and/or implementation of cellular markers have been established (16–18). Microbial discrimination by the host may be of particular importance for filter-feeding metazoans, such as corals, bivalves, or sponges, who in addition to discriminating between symbionts to retain and pathogens to eliminate, may have to recognize bacteria as food items.

The early branching phylum Porifera is an evolutionary and ecologically relevant group to study the role of phagocytosis in animal-microbe interactions. Sponges have no gut and the same phagocytic cell may retain a function in defense and nutrition (2). As benthic filter feeders, sponges constantly remove bacteria from the surrounding water as food items (19–21). At the same time, sponges harbor distinct microbial communities inside their mesohyl at up to 2-3 orders of magnitude higher abundances than in the surrounding seawater (22–24). These symbionts support the sponge metabolism through various processes, including carbon and nitrogen cycling as well as the synthesis of vitamins and secondary metabolites (e.g., (25–30). Recognizing and differentiating between food, symbiotic and/or pathogenic microbes is therefore paramount for these filter feeders. First observations of a reduced uptake of symbionts compared to seawater bacteria and the involvement of NLR-like immune receptors indicate such bacterial discrimination in sponges (31–33). Additional genomic studies further suggest that symbionts may avoid phagocytosis due to an enrichment of eukaryote-like proteins (e.g., ankyrins) presented on their surface (34–36), whereas seawater bacteria are taken up and digested. These proteins did promote bacteria persistence in cellular models, aiding the infection and survival of symbionts and pathogens in several eukaryotes, including amoeba, murine macrophages, and humans (37–39) by silencing innate immune responses. However, direct evidence if, and to what extent sponges facilitate bacterial discrimination on the cellular level is largely lacking.

A recently developed *in-vivo* phagocytic assay applying cell dissociation followed by fluorescence-activated cell sorting (FACS) in the sponge *Halichondria panicea* may now help to bridge this gap (40). In the present study, this assay was used to assess whether the sponge *H. panicea* can differentiate between two vibrio strains: Hal 281 isolated from *H. panicea* and is therefore defined as native and NJ 1 isolated from *Nematostella vectensis* and is hereafter defined as foreign. The relative abundance of sponge phagocytic cells (i.e., with incorporated fluorescently labeled *Vibrio*) was quantified by FACS. Phagocytic active cells were additionally inspected using fluorescence microscopy to determine their size and morphological type. A corroborative proteomics analysis was performed to investigate underlying molecular mechanisms of cellular recognition, phagocytosis, and immunity upon microbial exposure. Thereto, the total number of differentially abundant proteins responding to bacterial exposure after 30 and 60 min were analyzed.

## Materials and methods

### Sponge collection

Specimens of the sponge *H. panicea* (Pallas, 1766) were collected at the Kiel fjord (54.329659 N, 10.149104 E; Kiel, Germany) at 1 m water depth in August 2023, cleaned from epibionts, trimmed to approximately equally-sized fragments (volume: 32.3 ± 1.8 mL and wet weight: 5.9 ± 1.7 g [average ± S.D.]) containing 2-3 oscula, and left to heal and recover from collection on an *in-situ* nursery at the collection site for 10 days (41). On the day of the experiment, individuals were brought to the climate-controlled aquarium facilities of GEOMAR Helmholtz Centre for Ocean Research (Kiel, Germany), placed in a semi-flow through aquarium system supplied with natural seawater pumped from the collection site, and left to acclimatize for 2 h at 18°C room temperature, 17°C water temperature, and a salinity of 16-17 PSU, closely resembling environmental conditions at the nursery.

### Bacteria preparation for the in-vivo phagocytosis assay

The *Vibrio* isolates for the assay included the native sponge-associated isolate Hal 281 (isolated from *H. panicea*) and the foreign non-sponge-associated isolate NJ 1 (isolated from the sea anemone *Nematostella vectensis*). The similarity of 16S rRNA gene sequences between the two isolates was 95.62% (Table 1), and neighbor-joining phylogenetic analysis showed they belong to different clusters (Fig. S1A). *Vibrio* cultures were freshly grown in 100 mL liquid marine broth at 120 rpm, 25°C for 48 hours before the day of the experiment. The culture concentrations were estimated by OD_600_ and subsequently confirmed by flow cytometry measurements. The bacteria pellet was recovered by centrifugation (4000 x g for 10 min), resuspended in filtered artificial seawater (FASW), and fluorescently stained with TAMRA™ (Thermo Fisher Scientific, C1171) the same day of the experiment (5 μM final concentration, for 90 min in the dark, at room temperature). After the excess dye was washed off by centrifugation (4000 x g for 10 min), the bacteria were resuspended in FASW, and positive staining was confirmed with fluorescence microscopy (Fig. S1B). The *Vibrio* stocks were kept at 4°C until the experiment took place.

**Table 1.** Molecular and morphological comparison between the native and foreign *Vibrio* isolates Hal 281 and NJ 1, respectively. Molecular analysis is based on 16S rRNA gene sequences. The two closest type strains, based on blastn analysis on 16S rRNA gene are reported. The morphological features are reported as average ± standard deviation. GenBank accession numbers are given in parentheses.

| Isolate | Host | Sequence length (bp) and GenBank accession number | Closest type strains | Similarity (%) | Average ± SD cell length (μm) | Average ± SD cell width (μm) |
| --- | --- | --- | --- | --- | --- | --- |
| Hal 281 | *Halichondria panicea* | 1517 (MT406665) | *Vibrio atlanticus* LMG 24300 (EF599163)/<br>*Vibrio cyclitrophicus* NBRC 107756 (AB682659) | 99.47/<br>99.80 | 1.98 ± 0.25 | 0.90 ± 0.08 |
| NJ 1 | *Nematostella vectensis* (anemone) | 1455 (PQ455196) | *Vibrio diazotrophicus* NBRC 103148 (BBJY01000042)/<br>*Vibrio plantisponsor* MSSRF60 (GQ352641) | 99.40/<br>99.37 | 2.39 ± 0.86 | 0.31 ± 0.14 |

### Phagocytosis assay

We followed a similar methodology and experimental approach as described in (40), a brief description of the method follows. Individual specimens of *H. panicea* were placed in incubators filled with unfiltered seawater (natural seawater bacteria concentration: ∼ 10^5^ cells mL^-1^). Incubators with sponges were randomly assigned to one of the following treatments and incubated for either 30 or 60 min: Addition of native *Vibrio* (Hal 281, n=5), addition of foreign *Vibrio* (NJ 1, n= 5) or seawater control with no *Vibrio* addition (control, n = 5), (n: Biological replicates per treatment and per time point). The target concentration for the *Vibrio* treatments was 10^5^-10^6^ vibrios mL^-1^. Water samples for flow cytometry were taken through the incubation period to assess bacterial uptake by the sponge (Text S1, Table S1, and Fig. S2).

### Sponge cell dissociation

The entire sponge individuals were recovered at the end of each incubation and used to extract sponge cell fraction by a physical dissociation via differential centrifugation, as described in (40), (protocol adjusted after (32,42)). Briefly, the entire sponge fragments were rinsed with sterile, ice-cold Ca- and Mg-free artificial seawater (CMFASW (43)), cut into small fragments, and incubated for 15 min in ice-cold CMFASW with EDTA (25 mM) on a shaker. Samples were filtered through a cell strainer (40 µm) and the homogenate was centrifuged at 500 x g, for 5 min at 4°C. The supernatant was discarded, and the sponge cell pellet was resuspended in 4 mL of ice-cold CMFASW. Aliquots (1 mL) of the cell suspension were either sampled for proteomics analysis (see below) or fixed with PFA (final concentration 4%) overnight for estimating phagocytosis using FACS.

### Estimation of *Vibrio* phagocytosis by sponge cells

Phagocytosis of the *Vibrio* isolates by sponge cells was estimated using fluorescence-activated cell sorting (FACS) following the protocol by (40). Shortly, the fixative was washed off the cells by centrifugation (500 x g, for 5 min at 4°C) and the pellet was resuspended in 1 mL of ice-cold CMFASW. The concentration of the dissociated sponge cells was adjusted to approx. 3 x 10^7^ cells mL^-1^. Prior to the analysis, sponge cells were filtered again through a cell strainer (40 µm), and their nuclei stained with DAPI (final concentration 0.7 ng uL^-1^) to detect the “bulk” sponge cells population based on the dye fluorescence (355 nm UV laser and filter 448/59 nm). FACS analysis was performed on a MoFlo Astrios EQ® cell sorter (Beckman Coulter) using the Summit software (v6.3.1). Each sample was run five times and a total of 500k events were recorded for each technical replicate. The side scatter (SSC), DAPI, and the TAMRA-stained bacteria fluorescence (laser 561 nm and filter 692/75 nm) were used to identify and quantify the sponge cells that had incorporated the *Vibrio* isolates (40). The control sponges (i.e., individuals incubated without isolates) were used to correct for events corresponding to natural auto-fluorescence in the cells (as in (40)). We define sponge phagocytic cells as those that had incorporated the fluorescent vibrios during the assay, whereas non-phagocytic cells as those without incorporated vibrios (i.e., according to the presence or lack of fluorescence signal in the TAMRA channel, respectively). The relative (%) phagocytic and non-phagocytic cell fraction was estimated in relation to the total number of events from these two gates (for details see (40)). To test the effect of the *Vibrio* type (i.e., native *vs*. foreign) and of incubation time (i.e., 30 *vs.* 60 min), a two-way ANOVA analysis was performed (significance was determined at α = 0.05). Statistical analysis was performed in R-studio (V4.2.1; Rstudio Team 2022) by fitting an analysis of variance model (aov () function).

### Determination of phagocytic sponge cells using fluorescence microscopy

Dissociated sponge cell suspensions of three random individuals per treatment were sub-sampled and inspected by fluorescence microscopy to determine the cell types involved in phagocytosis of native and foreign vibrios. A total of 30 phagocytic cells per individual were counted (30 phagocytic cells x 3 replicates = 90 cells per treatment per time point), and the size of the cells as well as the number of TAMRA-stained *Vibrio* cells incorporated per cell was recorded. Cells were mounted on microscopy slides using ROTI mount FlourCare DAPI and examined under an inverted fluorescence microscope equipped with a camera (Axio Observer Z1 with Axiocam 506 and HXP-120 light; Zeiss), at a total magnification of 100x, using the filters 49 DAPI (335-383 nm for sponge nuclei) and 43 HE DsRed (538-562 nm for TAMRA-stained *Vibrio sp.*). ZEN Blue Edition software (Zeiss) was used for acquiring and editing pictures. Sponge cells were classified into four categories (Fig. 1A): (1) small-sized flagellated cells (3-6 µm Ø), (2) small-sized cells with no visible flagella (3-6 µm Ø), (3) medium-sized cells (7-10 µm Ø), and (4) big cells (> 10 µm Ø). As sponge cells change their typical morphology due to the dissociation process, we could only classify the cells based on their size and the presence/absence of flagellum. We further estimated the number of TAMRA-stained *Vibrio* cells that were incorporated per observed sponge phagocytic cell. The number of vibrios per sponge cell was divided into four categories (Fig. 1B): 1 vibrio, 2 vibrios, 3-5 vibrios, > 5 vibrios. One-way ANOVAs were run per treatment condition (i.e., per *Vibrio* type and per incubation time separately) to test if the cell types involved in the bacteria incorporation, as well as the number of *Vibrio* cells incorporated per phagocytic cell differ between the *Vibrio* isolates presented to *H. panicea*. Additionally, a PERMANOVA ((adonis2 () function), package vegan in R-studio) was performed in the combined data set to test if the distribution of phagocytic cell types as well as the number of vibrios incorporated per cell changed between *Vibrio* isolates and over time.

**Fig. 1.**
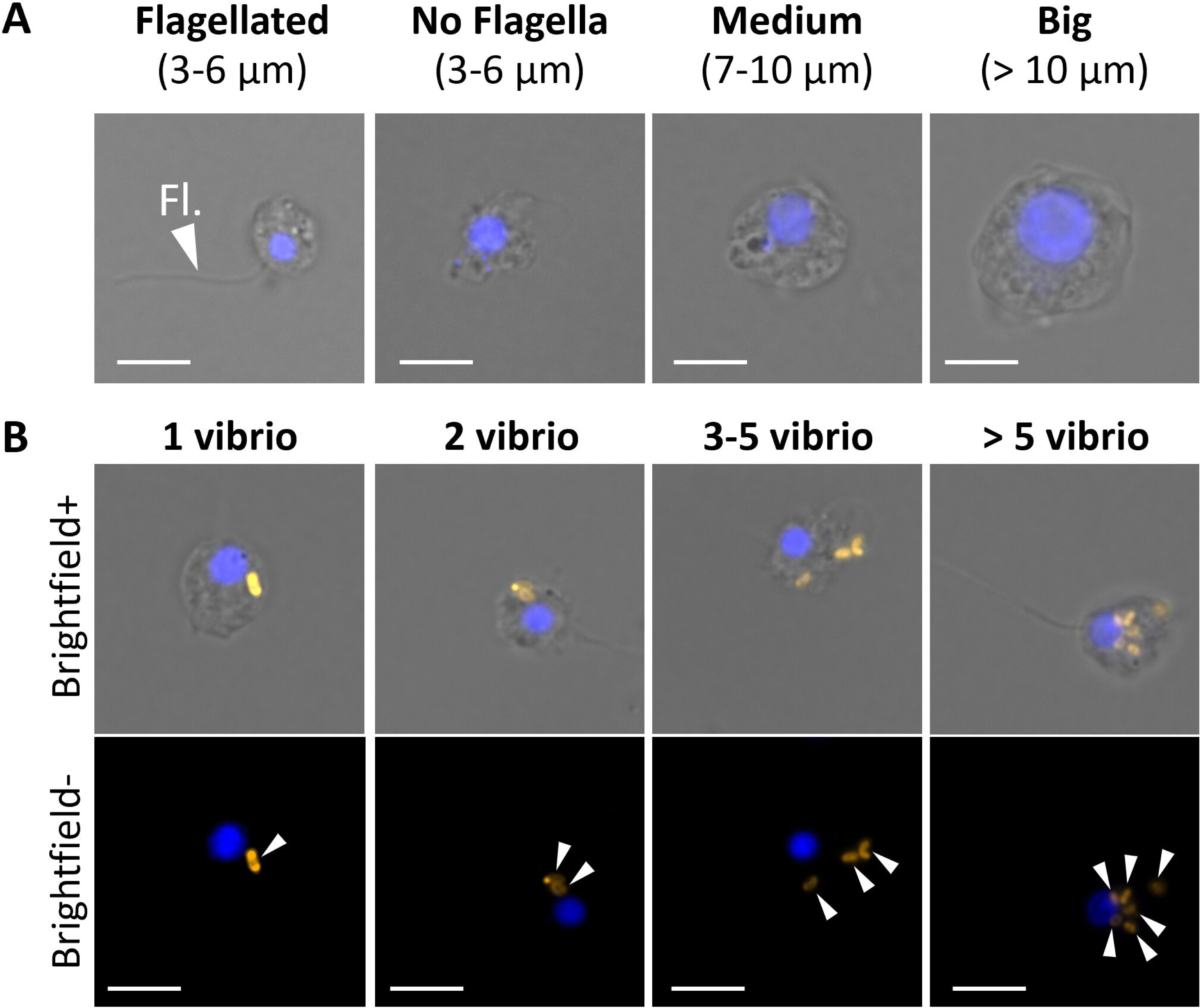
Phagocytic cell types of *H. panicea* individuals exposed to *Vibrio* isolates based on fluorescence microscopy. Representative fluorescent microscopy pictures showing (A) different phagocytic cell categories and (B) different amounts of *Vibrio* cells incorporated into sponge cells. Scale bars: 5 µm. Sponge cell nuclei (blue) stained with DAPI. Fl: Flagella. Arrowheads: TAMRA-stained *Vibrio*.

### Identification of differentially abundant proteins (DAPs) using proteomic analysis

Aliquots (1 mL) from the dissociated sponge cell suspensions were washed by centrifugation (500 x g, for 5 min at 4°C), the supernatant was discarded, and the cell pellet was resuspended in 1 mL of lysis buffer (5 M urea, 1% Triton x-100, 5 mmol L^-1^ Dithiothreitol, 1x cOmplete™ mini EDTA-free, and 50 mM Tris; pH 7.5-8). Samples were thoroughly vortexed for 30 s and incubated in a shaker (1500 rpm) for 40 min at 37°C. Throughout this step, samples were vortexed every 10 min for 30 s. Subsequently, they were placed in an ultrasonic bath for 5 min. Debris was pelleted by centrifugation (20,000 x g, for 30 min at 10°C), and the supernatant was transferred into a new tube and stored at -80°C until further processing. Protein concentrations were determined using the Pierce^TM^ BCA Protein assay kit (Thermo) according to manufacturer’s instructions. 50 μg protein of each sample was processed further according to the SP3 protocol (44) as follows. Proteins were reduced by addition of Dithiothreitol (DTT; 10 mM final concentration) and mixing at 1000 rpm, 56°C for 30 min. Alkylation was performed by addition of chloroacetamide (50 mM final concentration) and mixing at 1000 rpm, 25°C for 20 min. 50 μL of a mixture of hydrophobic and hydrophilic SP3 beads (20 μg μl^-1^) were added, reaching a 20:1 (beads:protein) ratio. Binding to beads was induced by adding ethanol to 54% and then mixing at 1500 rpm, 25°C for 15 min. Beads were washed 3 times, each washing step consisted of (1) centrifugation (21,000 g, 25°C for 2 min), (2) binding beads to a magnet and discarding the supernatant, (3) adding 400 μL of 80% Ethanol and ultrasonicating until beads disperse, and (4) finally vortexing for 10 s. The remaining supernatant was discarded after the last washing step, and beads were reconstituted in 100 μL of Triethyl ammonium bicarbonate (100 mM). Proteins were digested at 37°C overnight with 0.7 μg Trypsin/Lys-C mix (Promega). Afterwards, supernatant-containing peptides were vacuum dried then reconstituted in 100 μL 3% Acetonitrile (ACN) and 0.1% Trifluoroacetic acid (TFA). An additional desalting step was performed using Pierce™ C18 100 μL pipette tips as follows. Tips were washed twice with 100 μL (50% ACN) and equilibrated twice with 100 μL (3% ACN, 0.1% TFA). Peptides were loaded by pipetting up and down 10x and then washed 2x with 100 μL (3% ACN, 0.1% TFA). A first peptides elution was done with 100 μL 50% ACN and 0,1% TFA, and a final elution with 50 μL 70% ACN, 0.1% TFA. Eluted peptides were vacuum dried, reconstituted in 3% ACN, 0.1% TFA, and analyzed by liquid chromatography-mass spectrometry (LC-MS). Chromatographic separation was performed on a Dionex U3000 nano HPLC system equipped with an Acclaim pepmap100 C18 column (2 μm particle size, 75 μm × 500 mm) coupled online to a mass spectrometer. The eluents used were: Eluent A: 0.05% Formic acid (FA), eluent B: 80% ACN + 0.04% FA. The separation was performed over a programmed 132 min run. Initial chromatographic conditions were 4% B for 2 min followed by linear gradients from 4% to 50% B over 100 min, then 50 to 90% B over 5 min, and 10 min at 90% B. Subsequently, an inter-run equilibration of the column was achieved by 15 min at 4% B. A constant flow rate of 300 nL min^-1^ was employed. Two technical replicates of each sample were analyzed, and wash runs were performed between samples. Data acquisition following separation was performed on a QExactive Plus (Thermo). Full scan MS spectra were acquired (300-1300 m z^-1^, resolution 70,000) and subsequent data-dependent MS/MS scans were collected for the 15 most intense ions (Top15) via HCD activation at NCE 27.5 (resolution 17,500, isolation window 3.0 m z^-1^). Dynamic exclusion (20 s duration) and a lock mass (445.120025) was enabled.

MS raw data were analyzed against *H. panicea* predicted proteome (268,992 sequences; from (45)) and known contaminants (cRAP) using CHIMERYS identification search engine linked to Proteome Discoverer^TM^ 3.0 (Thermo). The raw data files can be found in the ProteomeXchange repository (identifier PXD-TBD) (46). Enzyme was set to Trypsin with two missed cleavages tolerance. 20 ppm fragment ion mass errors were tolerated. Carbamidomethylation of cysteine was set as a fixed modification and oxidation of methionine (M) was tolerated as a variable modification. Strict parsimony criteria were applied, filtering peptides and proteins at a 1% false discovery rate (FDR). Label-free quantification method based on the intensities of the precursor ions was used. Proteins were filtered to have “High” FDR combined confidence and at least two identified peptides. Data was further analyzed by Excel and Perseus v 1.6.15.0 (47). Protein intensities were averaged for technical replicates and then normalized by median-based normalization. Log2 transformed intensities were grouped into six groups depending on the *Vibrio* treatment and incubation time (each with four biological replicates) and filtered to contain four values in at least one group. Missing values were imputed from a normal distribution separately for each replicate (Width 0.3, Downshift 1.8). Statistical analysis was done using ANOVA, permutation-based FDR of 0.05 or 0.01, as indicated. Furthermore, we performed a direct pathway analysis (DirectPa R package) (48) on the quantified proteins based on the fold changes of each treatment relative to the control to better understand the sponge cells’ protein dynamics upon exposure to each *Vibrio* treatment.

### Functional annotation of proteins

Identified protein sequences were annotated to Uniprot identifiers of *Amphimedon queenslandica* proteome (Uniprot UP000007879_444682) and to KO by BlastKOALA using the Eukaryotes KEGG gene database (49). The best Blastp match with *A. queenslandica* was used as input in the DAVID web server (knowledgebase v2024q1) (50,51) to perform Gene Ontology (GO) functional annotation and enrichment analysis of the significantly differentially abundant proteins (DAPs). GO terms and fold enrichment values were analyzed in REVIGO (v1.8.1;(52)) to reduce redundancy and group them based on semantic similarity (SimRel).

The groups of proteins obtained from the directional analysis were further characterized to identify proteins likely to be involved in phagocytic, bacterial infection, and/or immunity pathways based on KEGG annotation. KEGG pathways were reconstructed with the KEGG Mapper Reconstruct web tool (v5) (53,54). The KEGG identifier numbers from the BlastKOALA annotation were used as input for reconstructing the pathways. KEGG pathways used for the protein classification included endocytosis (map04144), lysosome (map04142), and phagosome (map04145), bacterial infection (map05110, map05120, map05130, map05131, map05132, map05133, map05134, map05135, map05152) and Immunity (map04623, map04625, map04657, map04611, map04612, amp04613, map04620, map04621, map04622, map04624, map04666).

## Results

### Phagocytic active cell fraction after exposure to native and foreign *Vibrio* isolates

The average (± SD) number of sponge cells recovered after dissociation was approx. 4.89 x 10^7^ ± 3.00 x 10^7^ cells mL^-1^. The population of phagocytic cells of the *H. panicea* individuals incubated with the *Vibrio* isolates was clearly distinct from the corresponding gate in the control sponges (i.e., individuals incubated without isolates.) (Fig. 2A-C). In the 30 min assays, the percentage of phagocytic cells was on average 28.9 ± 2.6 % for Hal 281 and 28.2 ± 3.9 % for NJ 1 (Fig. 2D) with no significant difference between the isolates (one-way ANOVA, F = 6.1, *p* = 0.99; df = 3). In the 60 min assays, the average percentage of phagocytic sponge cells was 38.3 ± 3.8 % for Hal 281 and 33.6 ± 6.0 % for NJ 1. These percentages represent on average, a 9.3% and 5.4% increase for Hal 281 and NJ 1, respectively, compared to the estimates for the 30 min assays (Fig. 2D). There was no effect of the *Vibrio* isolate on the phagocytic activity of *H. panicea* cells (two-way ANOVA, F = 1.9, *p* = 0.19; df = 1; Fig. 2D), whereas incubation time significantly increased the number of bacterial incorporated into the sponge cells (two-way ANOVA, F = 15.9, *p* = 0.001; df = 1; Fig. 2D).

**Fig. 2.**
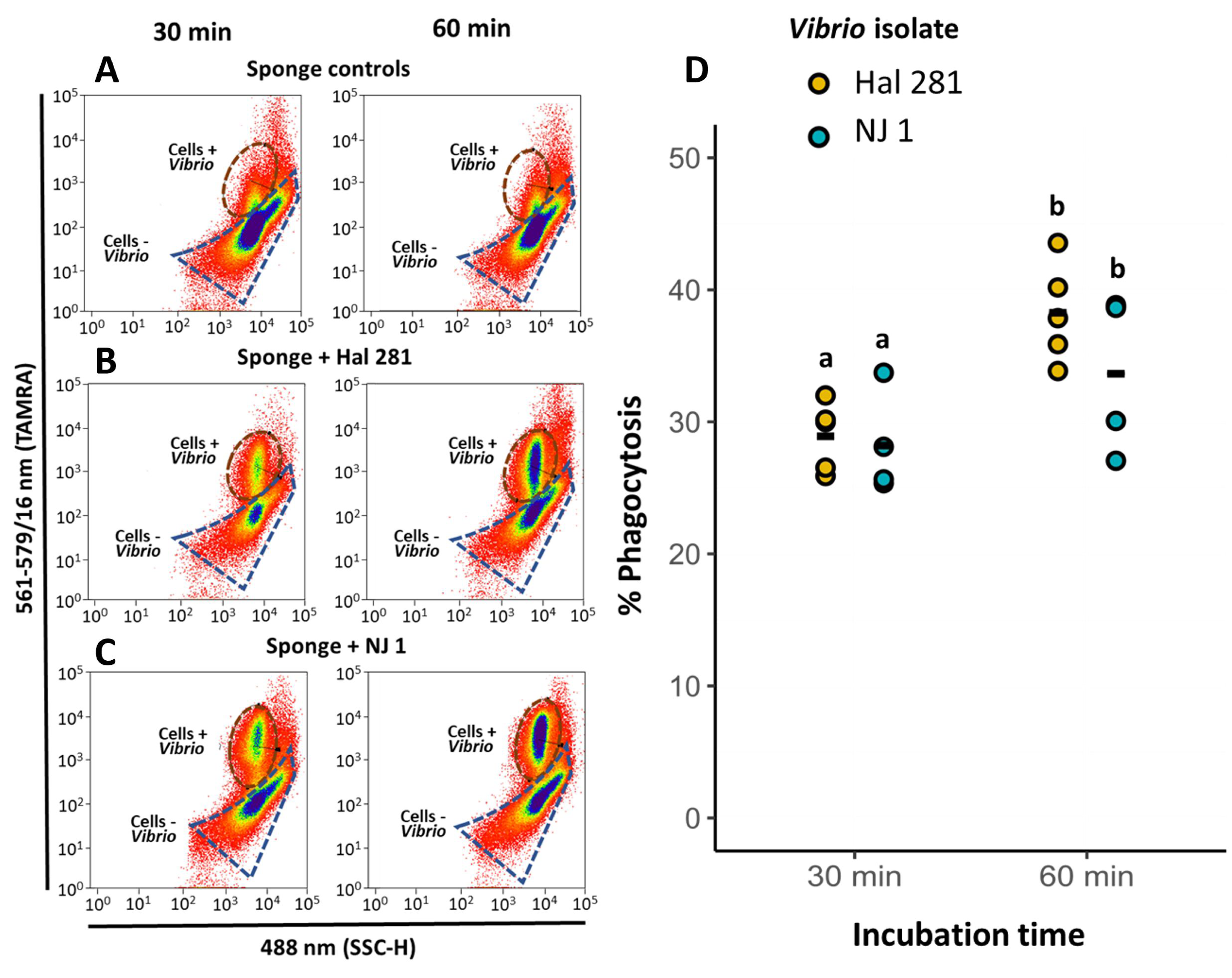
Fluorescence-activated cell sorting (FACS) analyses estimates of phagocytic cells. Representative cytograms for (A) sponge controls (incubated without *Vibrio* addition) and sponges incubated with the (B) native Hal 281 and (C) foreign NJ 1 *Vibrio* isolates, for 30 min (left) and 60 min (right). Brown and blue dashed outlines: Gates of phagocytic cells with (+) or without (–) incorporation of *Vibrio*, respectively. A total of 500k events were recorded in each case. (D) Estimates of phagocytic active sponge cells. The relative number (%) of cells phagocytizing *Vibrio* cells (+ TAMRA signal) to the total of sponge cells (DAPI-stained, SSC-H signal) was estimated via FACS analysis (whole data set in Table S2). Bold line: Average for the 4-5 biological replicates. Treatments marked with different letters are significantly different at ɑ=0.05.

### Distribution of *Vibrio* cells in different phagocytic cell types

Fluorescence microscopy observations of phagocytic cells (90 cells per treatment per time point) showed that *Vibrio* phagocytosis was performed by sponge cells with different sizes and morphologies (Fig. 1). In both *Vibrio* treatments, approx. 88 and 96% of the total phagocytic cells observed in the 30 min assays comprised small-flagellated cells (53-55%) and cells with no visible flagella (34-40%) (3-6 µm Ø. Fig. 3, Fig. S3 and Table S3). Medium (7-10 µm Ø) and big cells (> 10 µm Ø) only represented approx. 4-12% (i.e., 4-11 cells) of the recorded cells (one-way ANOVA, F = 18.6, *p* < 0.001, df = 7; Fig. S3). After the 60 min assays, the small-flagellated cells continued to be significantly higher than the other cell types (one-way ANOVA, F = 28.3, *p* < 0.001, df = 7; Fig. S3). When combining the two data sets, a significant interaction between time (30 *vs.* 60 min) and *Vibrio* isolate (Hal 281 *vs.* NJ 1) on the distribution of cell types involved in phagocytosis was revealed explaining 24% of the variation in the data (PERMANOVA, F = 7.8, *p* = 0.003, df = 1; Fig. 3 and Table S4A). Thus, the effect of time on phagocytic cell type distribution differed between the two *Vibrio* isolates. The percentage of flagellated cells and non-flagellated cells appeared to decrease from 30 to 60 min for both isolates, yet this decline was not significant (t-test flagellated: t = 0.61 and - 1.39, *p* = 0.60 and 0.30, df = 2; non-flagellated: t = 0.49 and 2.36, *p* = 0.67 and 0.14, df = 2, Hal 281 and NJ 1, respectively). Similarly, the medium-size cells tended to increase after the 60 min incubation for both Hal 281 and NJ 1 treated sponges, but also not significantly (t-test, t = -1.66 and -2.77, *p* = 0.24 and 0.11, df = 2, respectively).

**Fig. 3.**
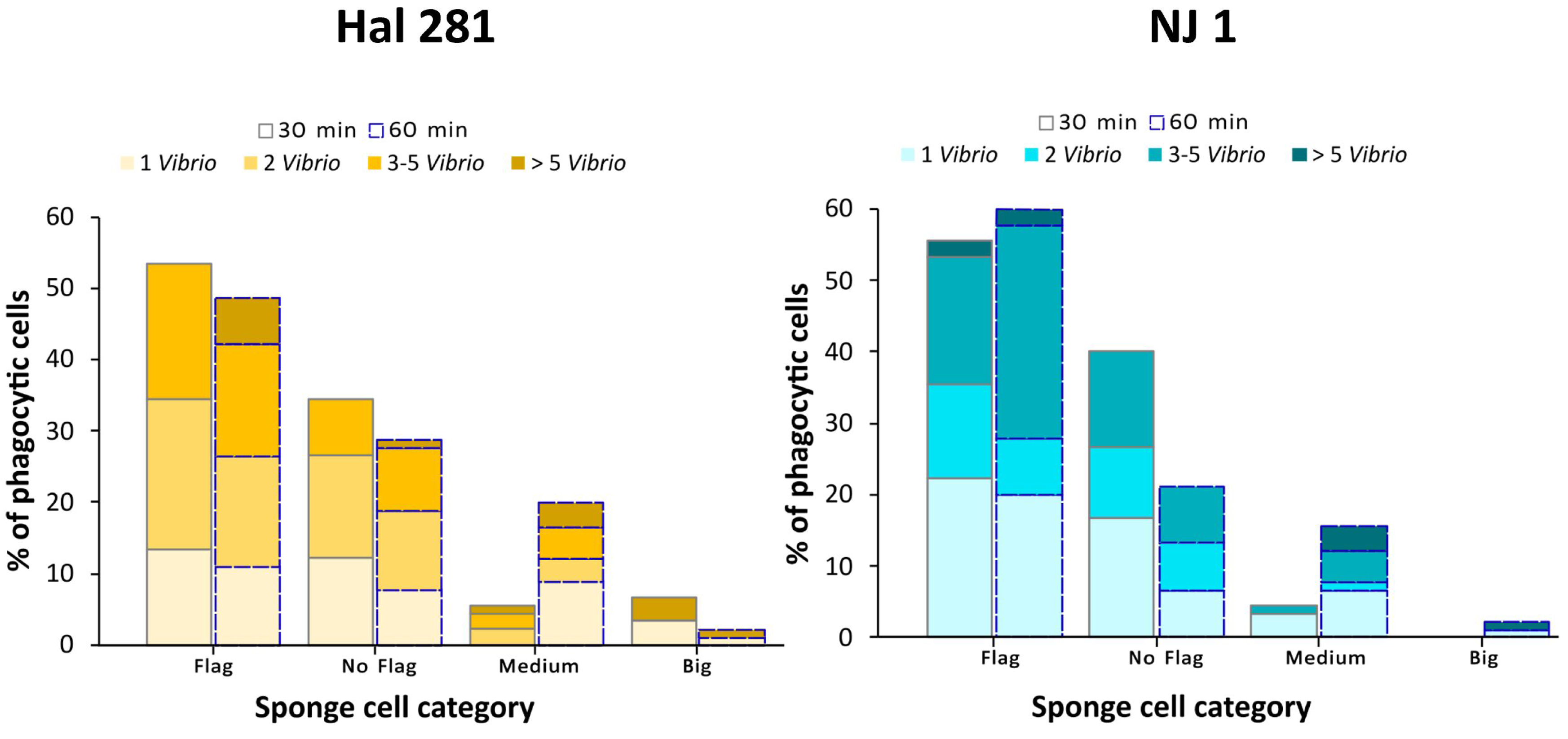
Distribution of *Vibrio* cells in the different phagocytic cell types after 30 min (gray outlined bars) and 60 min (dashed bars) incubations with the native isolate Hal 281 (left) and foreign isolate NJ 1 (right) based on microscopy cell counts. Flagellated and No flagellated: 3-6 µm Ø; Medium: 7-10 µm Ø; Big: > 10 µm Ø.

We further estimated the amount of *Vibrio* cells that were incorporated per phagocytic sponge cell (Fig. S4 and Table S3). Most sponge cells (88-98%) incorporated a maximum of 5 vibrio cells after both 30 and 60 min incubations (Fig. 3). The number of vibrios per sponge cell was not affected by incubation time but by *Vibrio* isolate (PERMANOVA, F = 2.9, *p* = 0.03, df = 1, Fig. 3 and Table S4B). The percentage of sponge cells with >3 vibrios was higher in sponges incubated with the foreign *Vibrio* NJ 1 than in sponges incubated with the native isolate Hal 281. Additionally, the distribution of *Vibrio* cells over the different types of phagocytic cells was significantly different between Hal 281 and NJ 1, and this effect of the *Vibrio* isolate explains 19% of the variation in the data (PERMANOVA, F = 2.5, *p* = 0.02, df = 1, Fig. 3 and Table S4C). The difference was especially evident in the flagellated cells of sponges incubated with NJ 1, which incorporated a higher number of vibrios than sponges incubated with Hal 281. Exposure time did not affect the distribution of vibrios among cell types.

### Differentially abundant protein profiles upon *Vibrio* and seawater bacteria exposure

In total, 4753 and 4893 proteins were quantified on sponge cell fractions after 30 and 60 min incubations, respectively (Fig. S5). The principal component analysis (PCA) revealed a distinct proteome signature of the NJ 1 treated sponges, which formed a separate cluster from control and Hal 281 in both 30 min (PERMANOVA, F = 1.78, p = 0.032, df = 2) and 60 min (PERMANOVA, F = 2.8, p = 0.001, df = 2) incubation datasets (Fig. S6A-B). Moreover, an effect of Hal 281 incubation on the sponge cells proteome was detected after 60 min incubations (Fig. S6A-B). Thus, 60 min exposure to the *Vibrio* isolates had the most prominent effect on the proteome of *H. panicea* cells.

We identified a total of 77 and 285 significant differentially abundant proteins (DAPs) between all the treatment groups after the 30 and 60 min incubations, respectively (ANOVA, permutation-based FDR = 0.05, Table S5, Fig. S6C-D). BlastKoala provided annotation for approx. 53-67 % of the total quantified DAPs at both time points (Fig. S6C-D. The full annotation report can be found in Table S5). At 30 min, a total of 55 DAPs were shared between the Hal 281 treated sponges and the control, whereas specimens exposed to NJ 1 only shared two DAPs with control sponges (Fig. S6C-D). After 60 min incubations, the total number of DAPs increased, but Hal 281 and control continued to share more DAPs than NJ 1 and control (117 and 41 proteins, respectively. Fig. S6C-D). Based on the abundance profile of DAPs within each sample, biological replicates were clustered according to treatment (Fig. 4A-B, upper dendrograms). The DAPs profile of sponges exposed to the native isolate (Hal 281) exhibited more similarity to the control at both time points, contrary to the foreign isolate (NJ 1) (Fig. 4A-B and Fig. S6A-B). Moreover, DAPs clustered into four and six different clusters based on their abundance profile similarities after 30 and 60 min, respectively (left dendrograms Fig. 4A-B).

**Fig. 4.**
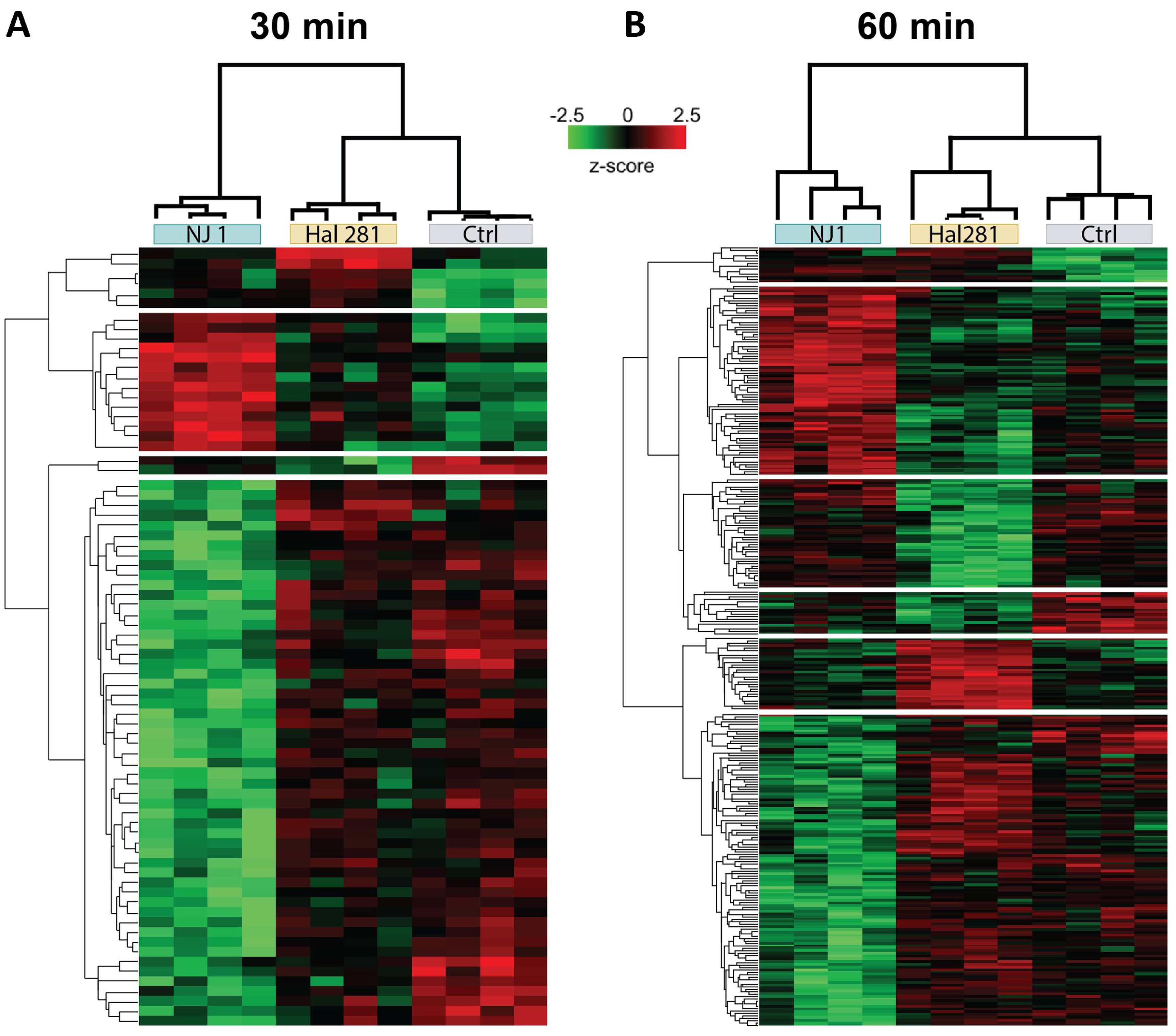
Heatmaps of differentially abundant proteins (DAPs) in *H. panicea* cells after 30 min (A) and 60 min (B) sponge incubations with the native and foreign *Vibrio* isolates Hal 281 and NJ 1, respectively. Sponges incubated without the addition of *Vibrio* isolates served as controls. Proteins were defined as differentially abundant relative to all the treatment groups with ANOVA permutated-based FDR p-value < 0.05, n = 4. Each row represents one protein, and each column one sponge sample. Dendrograms show protein clusters with similar abundance based on Euclidean distance and normalized z-score. Significant clusters are indicated by breaks in the heatmap.

### Protein-related functions of *Vibrio* exposure relative to seawater control

We further performed a direction analysis pathway (directPA) (48) on the quantified proteins based on the fold changes of each treatment relative to the control to better understand the protein dynamics elicited by the two *Vibrio* strains. According to this analysis, we defined four DAP categories: High or Low abundance in both *Vibrio* treatments (H-H and L-L, respectively), High abundance in Hal 281 but Low abundance in NJ 1 (H-L), and Low abundance in Hal 281 but High abundance in NJ 1 (L-H) (Fig. 5A-B, Table S6 and S7). The two last categories represent *Vibrio*-specific proteomic responses. At 30 min, 222 and 174 proteins were identified in the H-H group and the L-L group, respectively (Fig. 5A-B). In addition, the abundance of 24 proteins increased in response to Hal 281 but lowered in NJ 1 treated sponges (H-L) (Fig. 5A-B, Table S6). At 60 min, the number of proteins observed to be high (H-H) and low (L-L) in *Vibrio*-treated individuals decreased to 195 and 129 proteins, respectively (Fig. 5A-B, Table S7). Conversely, the proteins with different abundance in each specific *Vibrio* treatment increased to 68 and 66 proteins for the H-L and L-H groups, respectively (Fig. 5A-B). These results indicate an increase in *Vibrio-*specific changes in the proteome over time.

**Fig. 5.**
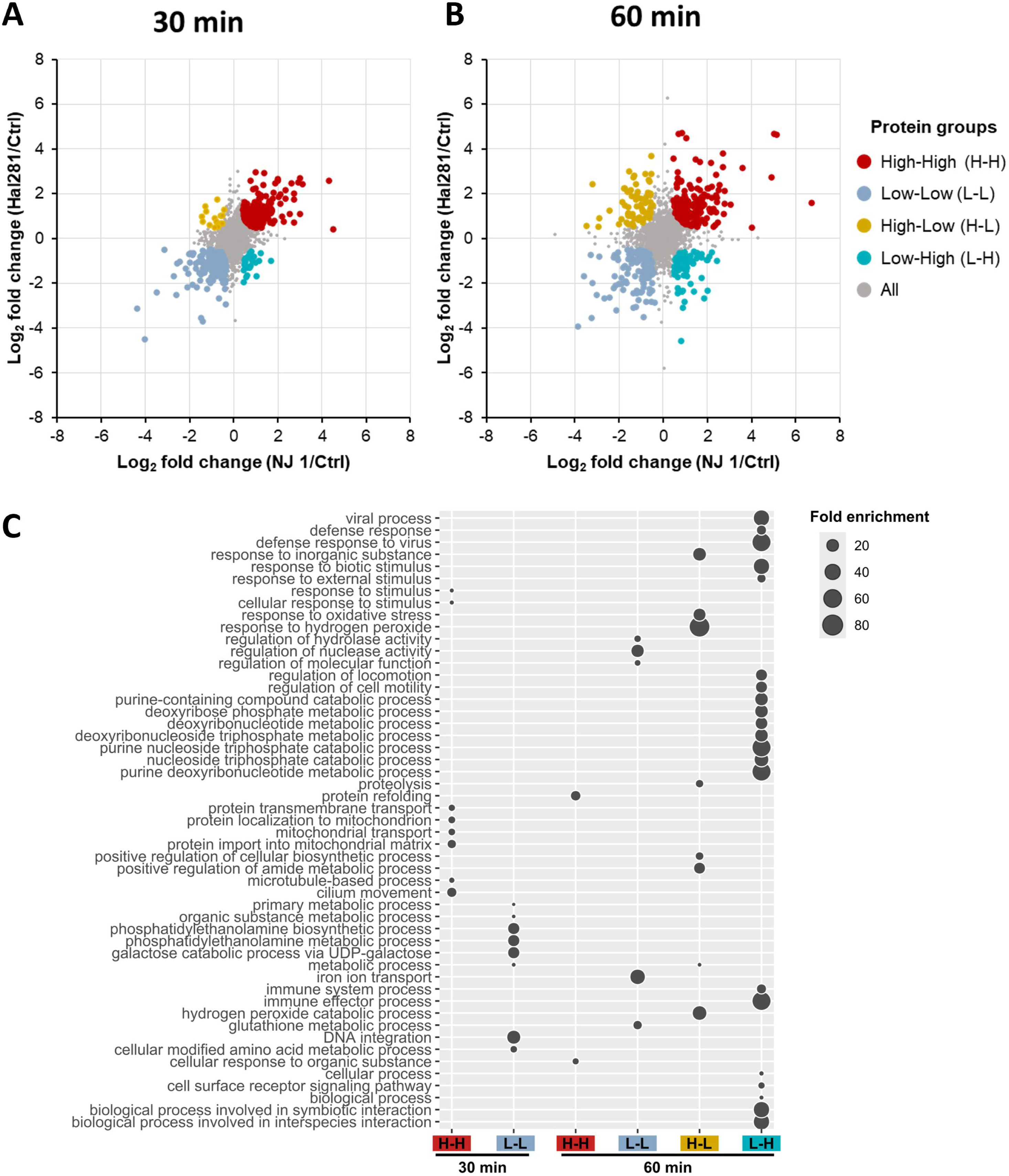
Directional pathway analysis of proteins identified after incubating *H. panicea* with a sponge-associated (Hal 281) and a non-sponge-associated (NJ 1) *Vibrio* isolate for (A) 30 min and (B) 60 min. Individuals incubated without the addition of *Vibrio* isolate served as control (n = 4 biological replicates per treatment). Proteins with higher or lower abundances in both treatments relative to the control (Ctrl) using protein fold changes and with a high probability (p-value < 0.01) are shown. (C) Predicted functions of differentially abundant proteins that were high or low in *H. panicea* cells after exposure of 30 min and 60 min to the native and foreign *Vibrio* isolates Hal 281 and NJ 1, respectively. Functions are based on GO annotations assigned by DAVID as biological processes. Functional descriptions correspond to REVIGO semantic clustering. Circle size indicates the fold enrichment value of each function. This bubble plot is equivalent to the tree maps generated by REVIGO (dispensability threshold < 0.5). Only groups with more than 30 proteins with GO annotations are shown in the plot. H-H and L-L: Proteins were high and low in response to both *Vibrio* isolates, respectively. H-L: Proteins were high in Hal 281 treated sponges but low in NJ 1 exposed individuals. L-H: Proteins were high in NJ 1 treated sponges but low in Hal 281 exposed individuals.

Gene Ontology (GO) enrichment analysis of proteins belonging to the four DAP categories revealed no significant enrichment (FDR corrected p-value < 0.05). Yet, we used the GO annotation (Table S8) to predict the functions in the DAP categories containing more than 30 proteins with GO annotation. After the 30 min incubation, *H. panicea* sponge response to both *Vibrio* isolates consisted mainly of an increased abundance (H-H category; 175 annotated DAPs) of proteins involved in processes related to protein import to mitochondria (GO:0006839, GO:0070585 and GO:0071806) and microtubule-based functions (GO:0003341 and GO:0007017), and a decrease (L-L; 119 annotated DAPs) of those associated with metabolic processes comprising phosphatidylethanolamine (GO:0046337 and GO:0006646) and galactose (GO:0033499) (Fig. 5C). The proteomic response of sponges after 60 min incubation with *Vibrio* showed an increased abundance (H-H; 130 annotated DAPs) in proteins related to protein refolding (GO:0042026) and cellular response to organic substances (GO:0071310), as well as a decreased abundance (L-L; 86 annotated DAPs) in proteins associated with regulation of molecular functions (GO:0065009, GO:0032069 and GO:0051336), glutathione metabolic process (GO:0006749) and iron transport (GO:0006826) (Fig. 5C). In addition, we could also predict the functions of proteins with *Vibrio*-specific responses: Hal 281 (H-L; 43 annotated DAPs) increased the abundance of proteins associated with oxidative stress (GO:0006979 and GO:0042542) and response to inorganic substances (GO:0010035), whereas NJ 1 (L-H; 46 annotated DAPs) increased proteins assigned to functions associated with a response to stimulus, purine deoxyribonucleotide metabolism (GO:0009151, GO:0009143, GO:0009146, GO:0009200, GO:0009262, GO:0019692, and GO:0072523), immune system (GO:0002252 and GO:0002376), and biological processes in interspecies interactions (GO:0008150, GO:0044419, and GO:0044403) (Fig. 5C).

### Proteins involved in phagocytosis, bacterial infection, and immunity upon *Vibrio* exposure

The groups of proteins obtained from the directional analysis were further characterized to identify proteins associated with phagocytic, bacterial infection, and/or immunity pathways based on KEGG annotation (see material and methods for details and Table S9). At 30 min, a total of 25 proteins were classified under these KEGG terms, and 16 of these exhibited high abundance profiles (H-H), whereas seven showed low abundances (L-L) in response to both *Vibrio* isolates, compared to the control treatment (Fig. 6A). Among the proteins in the H-H group, we detected endocytic (ko04144) and lysosome (ko04142) proteins, such as a heat shock protein 70 (HSPA1_6_8; K03283), a charged multivesicular body protein (CHMP1; K12197), and a vacuolar protein sorting-associated protein (VTA1; K12199). Among the proteins in the L-L group, we found a Ras-related protein (Rab-10; K07903), a cytohesin (CYTH; K18441), and two lysosomal-associated membrane proteins (LAMP1; K06528). Lysosomal proteins such as phospholipases (LYPLA3; K06129) were detected both within the H-H and L-L categories (Fig. 6A). Proteins predicted to be related to bacterial infection increased in abundance; for example, dehydrogenase (GAPDH; K00134), a ribosomal protein (RPS3; K02985), an associated death domain protein (FADD; K02373), and two exocyst complex components (EXOC5; K19984 and EXOC7; K07195) (Fig. 6A). In total, nine proteins were annotated as immune-related pathways, and seven of these proteins showed high abundance profiles (H-H), in response to both *Vibrio* isolates, compared to the control treatment (Fig. 6A). The immune-related proteins included for example a dual oxidase (DUOX; K13411), a proteasome activator subunit (PSM3; K06698), a member of the cytochrome P450 enzyme family (TBXAS1; K01832), and two histones (H2A; K11251) (Fig. 6A).

**Fig. 6.**
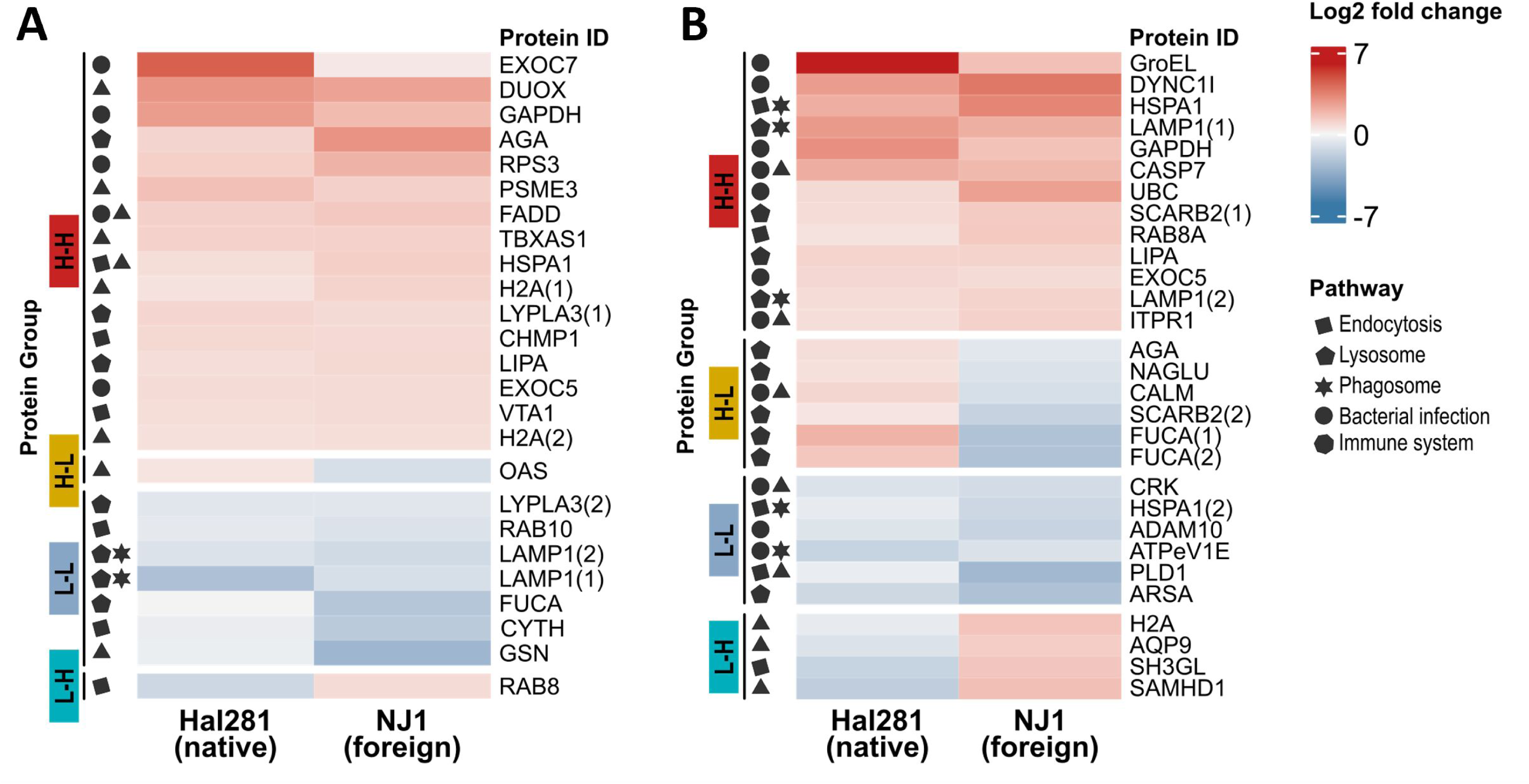
Predicted significantly differentially abundant proteins (DAPs) involved in pathways related to phagocytosis, bacterial infection, and immunity after (A) 30 min and (B) 60 min sponge incubations. Annotation is based on KEGG terms of the DAPs identified in *H. panicea* cells after exposure to the native and foreign *Vibrio* isolates Hal 281 and NJ 1, respectively. Color gradient: Protein abundance Log2 fold change relative to the control treatment. Each row represents one protein and each column one sponge sample. Significant protein clusters with similar abundance are indicated by breaks in the heatmaps. Numbers in brackets indicate different proteins with the same annotation. KEGG pathways included endocytosis (map04144), lysosome (map04142), and phagosome (map04145), bacterial infection (map05110, map05120, map05130, map05131, map05132, map05133, map05134, map05135, map05152) and Immunity (map04623, map04625, map04657, map04611, map04612, amp04613, map04620, map04621, map04622, map04624, map04666).

At 60 min, we found 29 proteins related to phagocytic-, bacterial infection-, and/or immune-related pathways (Fig. 6B). Both *Vibro* isolates continued to increase the abundance of the lysosomal proteins LAMP1 and LIPA (Fig. 6B) and further increased the abundance of a Ras-related protein (RAB8A; K07901), a lysosomal membrane protein predicted to have scavenger receptor activity (SCARB2; K12384), and a dynein protein related to microtubule movement (DYNC1l; K10415) (Fig. 6 B). In contrast, proteins related to phagocytosis, which decreased in abundance in *H. panicea* cells as a response to the *Vibrio* encounter (L-L), included a phospholipase D (PLD1; K01115), an ATPase (ATPeV1E; K02150), and an arylsulfatase (ARSA; K01134) (Fig. 6B). The number of proteins with opposite abundance regulation upon Hal 281 and NJ 1 (H-L and L-H) increased from one protein to up to six proteins from 30 min to 60 min. Among these proteins, five lysosomal proteins significantly increased after exposure to Hal 281 and decreased in sponges treated with NJ 1 (H-L; Fig. 6B). These included two α-fucosidases (FUCA; K01206), two glucosaminidases (AGA; K01444 and NAGLU; K01205), and a SCARB2 (K12384). Contrastingly, immune-related proteins, such as a histone (H2A; K11251), an aquaporin (AQP9; K09877), and a deoxynucleoside triphosphate triphosphohydrolase (SAMHD1), significantly increased after exposure to NJ 1 and decreased in sponges treated with Hal 281, compared to control individuals (Fig. 6B).

## Discussion

In the current study, we analyzed the cellular and proteomic response of *H. panicea* upon encounter with a native, sponge-isolated *Vibrio* (Hal 281) and a foreign, anemone-isolated *Vibrio* (NJ 1) after 30 and 60 min exposure using the combination of a recently developed *in-vivo* assay with proteomics analysis. The percentage of sponge cells engaged in phagocytic activity was similar between the tested isolates. Yet, the phagocytic response differed between Hal 281 and NJ 1, both in the sponge cell types involved in phagocytosis as well as the number of vibrios incorporated per cell. Sponges exposed to the native Hal 281 *Vibrio* showed a proteomic signature that was more similar to naïve (control) sponges than to foreign NJ 1-treated sponges. Further differences between the *Vibrio* isolates in their proteomic profiles pointed to an increased abundance of proteins related to lysosomal transport and digestive functions for the native Hal 281, and immune and inflammatory functions-related proteins for the foreign NJ 1.

### Choanocytes likely comprise the majority of phagocytic cells

Fluorescence-activated cell sorting revealed that more than 50% of the phagocytic active cells (i.e., with internalized vibrios) were in the size range of 5-6 µm in both *Vibrio* treatments and time points (Fig. 1, Fig. 3, and Table S3). Based on their (1) size range [*H. panicea* choanocytes: 3-7 µm (55,56)], (2) earlier reports of rapid incorporation of bacteria and small particles (≤ 2 µm) into choanocytes (e.g., (26,57)), and (3) the presence of a flagellum (58), we propose that these active phagocytic cells are choanocytes. We also observed cells in this size range without visible flagella (Fig. 1A). These cells may as well be choanocytes that either lost their flagellum during the cell dissociation process or the flagellum was not noticeable due to the limited resolution obtained by fluorescence microscopy, as previously described using scanning electron microscopy (59). An alternative explanation is that these, or at least a fraction of these small non-flagellated cells represent pinacocyte-like cells. Pinacocytes presumably capture large particles (> ostia diameter) by filopodial extensions and intracellularly digest them to prevent clogging of the sponge’s filtering system (60–62). The medium (7-10 µm) and big (>10 µm) phagocytic active cells (Fig. 1A) are most likely archaeocyte-like cells, given their bigger size and the presence of a large nucleus, which in some cases contained a distinctive nucleolus. Archaeocytes are amoeboid totipotent stem cells that move throughout the sponge mesohyl to participate in intracellular digestion and transport of food and particles between cells (63–65), but also neutralize foreign material as an immune defense system (2).

### Incorporation into phagocytic cells is *Vibrio* strain-specific

Cellular processing of the native (Hal 281) and the foreign (NJ 1) *Vibrio* isolates appeared to be largely similar with comparable abundances of phagocytic active cells (Fig. 2D). Also, the distribution of phagocytic active cell types showed a similar temporal pattern for both *Vibrio* isolates: The percentages of small flagellated and no-flagellated cells tended to decrease from the 30 to the 60 min time point, whereas those of medium and big cells appeared to increase. This trend may suggest that choanocyte-like cells initially incorporate vibrios and start to digest them (30 min time point; see general proteomic response to vibrios below). Once a saturation capacity is reached (e.g., five or more *Vibrio* cells, Fig. 3), predigested vibrios are translocated to archeocyte-like cells for further processing and transport within the mesohyl (60 min time point). Within this cellular processing, we however detected some strain-specific differences. Phagocytic cells (particularly those in the small size range, 3-6 µm) had more NJ 1 incorporated per cell than Hal 281 (Fig. 3). This strain-specific cellular response could be explained by a faster transfer of the native Hal 281 from choanocyte-like cells to archeocyte-like cells compared to the foreign NJ 1, as previously suggested for native *vs.* foreign bacteria in the sponge *Amphimedon queenslandica* (66). Alternatively, the initial incorporation of the foreign NJ 1 could be faster than that of the native Hal 281, or Hal 281 is evading initial digestion in choanocyte-like cells and is therefore transferred faster. All three options are not mutually exclusive and such microbe-specific differences in cellular processing appear to be widespread among marine invertebrates, including sea urchin larvae (67), deep-sea mussels (68), and sea anemones (16) (Table S10).

### Proteomic response is mainly driven by distinction between known and unknown bacteria

The proteomic profile of *H. panicea* samples incubated with Hal 281 was more similar to the control (i.e., exposure to natural seawater bacterial consortium) than to the foreign *Vibrio* treatment NJ 1 and this difference was more pronounced in the 60 min compared to the 30 min assays (Fig. S6). This pattern suggests a mechanism for specificity driven by the distinction between known (i.e., native Hal 281 and commonly encounter seawater bacteria) and unknown (i.e., foreign NJ 1) bacteria, rather than by bacterial type (i.e., *Vibrio vs*. seawater consortia). Hal 281 typically occurs in low abundances within *H. panicea* and can also be found in seawater samples collected close to sponges (69), making it more likely to be a commensal, rather than an obligatory symbiont. Since *H. panicea* is filtering approx. 1 L of seawater g dry weight^-1^ h^-1^ (70,71), it is expected to process millions of seawater bacteria per day, including *Vibrio* Hal 281. In contrast, the NJ 1 *Vibrio* isolate occurs in high abundance in juveniles of the sea anemone *N. vectensis* and is considered a native colonizer of this species (72). Since it is typically not found in *H. panicea* environment, it is unlikely that *H. panicea* has encountered it before. Our interpretation of specific discrimination between native or “known” microbes *vs*. foreign or “unknown” microbes implies a potential for immune memory in sponges that remains to be tested. While we interpret this distinction from the host’s perspective, potential mechanisms employed by each *Vibrio* to evade host recognition or degradation must also be considered. For example, the mechanisms for NJ 1 to colonize *N. vectensis* may be the ones that elicit an innate immune response in a different host, in this case, *H. panicea*. In both interpretations, commensal bacteria could still be partially digested but only the “unknown” or not co-evolved microbe will elicit an innate immune response.

### *Vibrio* encounter elicits a generalist mitochondrial protein-mediated response

When comparing the proteomic response of each *Vibrio* isolate to the respective control treatment (i.e., directional analysis), most differentially abundant proteins (DAPs) were similar between Hal 281 and NJ 1. These proteins appear to constitute a generalist response of *H. panicea* to *Vibrio* encounters, irrespective of strain. This generalist response features an increase in proteins related to mitochondrial transport, microtubule-based processes, protein refolding, as well as response to stimulus, and a concomitant decrease in proteins associated with metabolic processes, iron transport, and glutathione metabolism (Fig. 5C and Table S8). Some of these functions are further related to bacterial infection and/or immune-related pathways (based on KEGG annotation; Fig. 6 and Table S9), particularly mitochondria-mediated responses (73,74). Mitochondria are recognized to play important roles in sensing and responding to cellular and environmental signals by regulating cytosolic protein synthesis and gene expression to ensure cellular homeostasis (75–78). In *H. panicea* cells the abundance of a caspase (CASP7; K04397) and a cytochrome (TBXAS1; K01832) protein was high in response to both *Vibrio* isolates (H-H in Fig. 6) compared to the control treatment. Cytochrome proteins participate in cellular energy metabolism and caspase activation (79,80). Though caspases are commonly associated with apoptosis, these endoproteases are also involved in non-apoptotic processes such as cell proliferation, differentiation, and cytoskeletal reorganization (81,82). Thus, we speculate that following the internalization of *Vibrio* cells by *H. panicea*, a mitochondrial protein-mediated response is activated, potentially integrating with other downstream cellular processes.

### Native *Vibrio* elicits digestion, foreign *Vibrio* triggers an immune response

Differences in DAPs between the native (Hal 281) and the foreign (NJ 1) *Vibrio* isolates only became more evident after 60 min (i.e., proteins responding specifically to each isolate relative to the control; H-L and L-H Fig. 5C and Fig. 6B). Hal 281 elicited an increase of proteins related to oxidative stress, amide metabolism, response to inorganic substances, and proteolysis compared to control sponges (but less abundant in NJ 1 treated sponges. H-L Fig. 5C and Table S8). In fact, six out of the seven highly abundant proteins responding to the native *Vibrio* treatment were predicted to be lysosomal acid hydrolases (AGA, FUCA, NAGLU) and a lysosomal membrane protein (SCARB2) from the lysosomal pathway (map04142) (Fig. 6B and Table S9). This may suggest that after 60 min the native isolate (Hal 281) is already in the sponge’s phagolysosomes. After phagosome-lysosome fusion, reactive oxygen species (ROS) are produced to eliminate invasive microbes as part of the digestion process (7,83). Therefore, we hypothesize that the oxidative stress response observed after 60 min exposure to Hal 281 may indicate ROS production from the digestion of this *Vibrio* in the sponge phagolysosomes (Fig. 7). This digestion process likely corresponds to nutritional purposes. In contrast, the foreign *Vibrio* (NJ 1) caused an increase in DAPs related to immunity, purine metabolism, and interspecies interactions compared to control sponges (but less abundant in Hal 281 treated sponges. L-H Fig. 5C and Table S8). Purinergic signaling induces the activation of inflammatory response in immune cells such as macrophages, which results in a downstream release of caspases, interleukins, and cytokines upon exposure to LPS (84). The directional analysis and KEGG annotation further showed that three out of the four highly abundant proteins responding to the foreign *Vibrio* treatment were related to immune-related pathways (group L-H Fig. 6B). Two of these were annotated as nuclear proteins (SAMHD1 and H2A. Fig. 6B and Table S9), which could work as immune modulators (85,86) and even as immune effectors (87). The third highly abundant immune-related protein was an aquaporin (AQP9. Fig. 6B and Table S9), which together with H2A, is related to the formation of neutrophil extracellular trap (NET) proteins (88). NETs consist of cell material released during cell death that traps and kills pathogens either through antimicrobial proteins or by immobilizing microbes and facilitating their incorporation into phagocytic cells (85,89). We hypothesize that after initial ingestion, NJ 1 is recognized as a foreign agent and therefore as a potential threat, which elicits an inflammatory immune response resembling NETs (Fig. 7). These findings on the differential processing of native *vs.* foreign vibrios in *H. panicea* provide evidence of immune specificity in sponges.

**Fig. 7.**
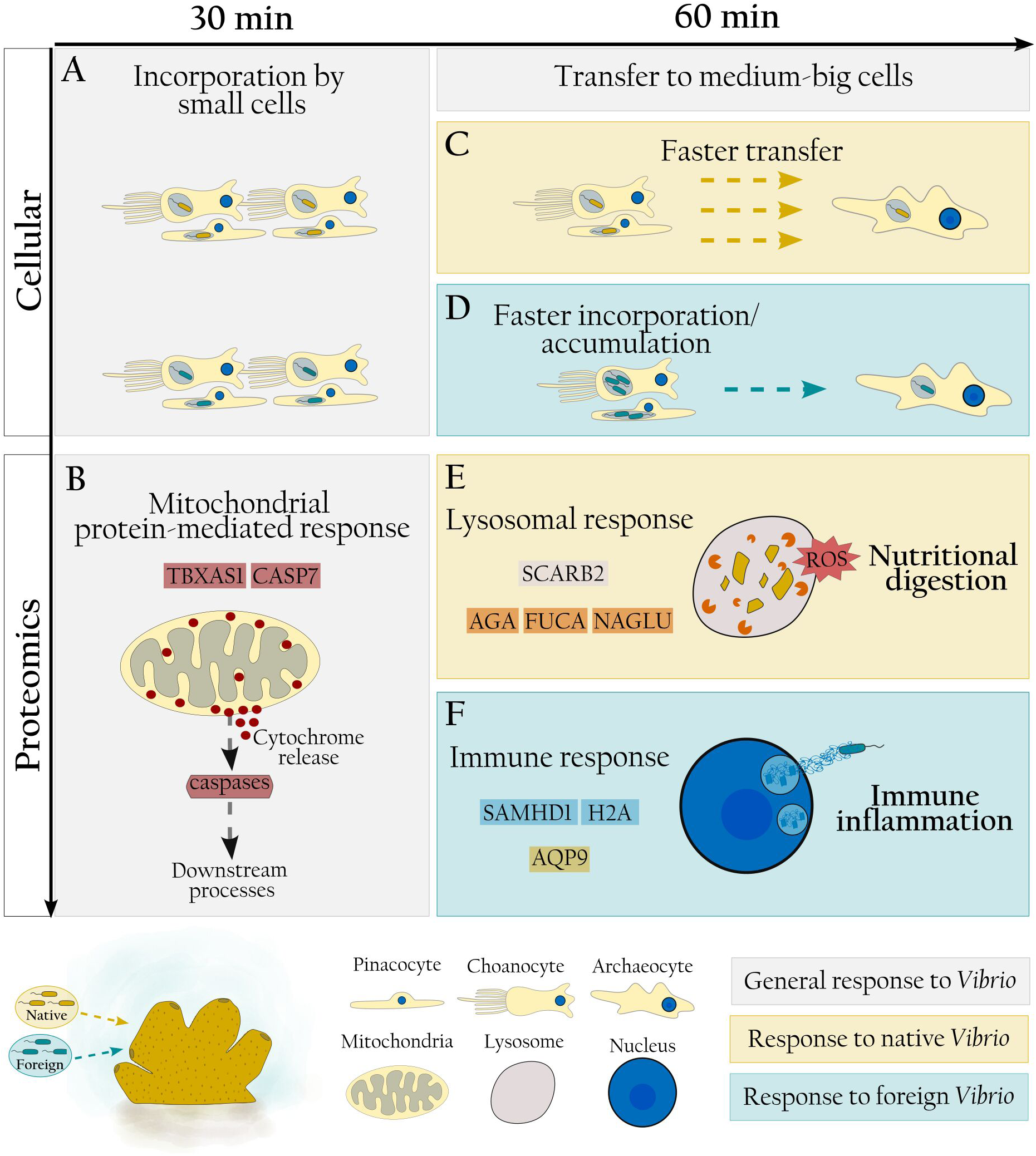
Proposed cellular and proteomic response to exposure to native and foreign vibrios in *H. panicea*. (A) Native and foreign vibrios are indiscriminately incorporated in small phagocytic cells upon encounter and (B) elicit a general mitochondrial protein-mediated response. (C) The native *Vibrio* strain is rapidly transferred to medium and big cells, whereas (D) the foreign *Vibrio* strain is accumulated to higher numbers and/or incorporated faster before transfer. (E) Increased abundance of proteins related to lysosomal activity and oxidative stress after 30 min suggests regular digestion of native vibrios. (F) Foreign vibrios appear to cause an increase in immune- and inflammatory-proteins after 30 min indicating a defense response against a potential threat.

### Proposed immune specificity in the processing of native and foreign vibrios

Based on combined cellular and proteomic results presented here, we propose that native and foreign vibrios are indiscriminately incorporated by choanocytes upon contact (Fig. 7A). Incorporated bacteria are first recognized as vibrios eliciting a mitochondrial protein-mediated response, involving cytochrome and caspase release, which integrates with other cellular functions operating downstream, such as energy metabolism, cell proliferation and differentiation, and cytoskeletal reorganization (Fig. 7B). In a second recognition step, vibrio strains are discriminated from one another: Native vibrios are considered food items and rapidly transferred to archaeocytes (Fig. 7C). There bacterial degradation is ultimately completed, as indicated by increased lysosomal activity and the occurrence of oxidative stress as part of the digestion process (Fig.7E). In contrast, foreign vibrios are accumulated in choanocytes and/or incorporated faster before being transferred to archaeocytes (Fig. 7D). Foreign vibrios may be considered a threat and trigger inflammatory and immune responses, such as the activation of immune modulators and/or effectors, and the formation of NETs to eliminate the invader (Fig. 7F). This differential recognition and processing of native and foreign vibrios highlights the presence of immune specificity mechanisms in *H. panicea*.

## Conclusion

The application of the *in-vivo* phagocytosis assay in the sponge *H. panicea* in combination with fluorescence microscopy and proteomics proved to be a powerful tool to elucidate the discrimination between native and foreign bacteria via phagocytosis, a conserved cellular mechanism from amoeba, to sponges, and humans. Our results suggest a clear discrimination between native and foreign vibrios in *H. panicea*, which occurs in two steps: (1) an early non-specific recognition and incorporation, discriminating vibrios from seawater bacteria and eliciting a mitochondria-mediated response (within 30 min). (2) A strain-specific discrimination (after 30 min) resulting in the digestion of the native strain *vs.* triggering an inflammatory immune response towards the foreign strain. This recognition and strain-differential processing of native and foreign microbes indicates immune specificity and may support the existence of immune memory in sponges.

## Data availability

The authors confirm that the data supporting the results and conclusions of this study are accessible within its supplementary information. If further data is required, it will be made available by the authors on request.

LC–MS raw data files have been deposited to the ProteomeXchange Consortium by the PRIDE partner repository with the dataset identifier PXD-TBD.

## Supplementary material

The supplementary material can be found online.

## Authors contributions

AMMG, LP, SF, and UH conceived the idea. AMMG and BM planned and conducted the experiments. KB performed the FACS analysis and AMMG the fluorescence microscopy inspections. MA and AT performed the proteomics analysis. The initial draft of the manuscript was written by AMMG. All authors contributed to improving the article and approved the submitted version.

## Acknowledgments

We are grateful to Nida Kaya for helpful discussions on the experimental design and for providing the isolates from *N. vectensis*. We further acknowledge Andrea Hethke for technical assistance in the lab, and Jutta Wiese and Tanja Rahn for their support with the sequencing and phylogenetic analysis of the bacteria isolates. We thank the International Max Planck Research School for Evolutionary Biology and the Collaborative Research Centre “Origin and Function of Metaorganisms” (CRC1182) for the supervision effort of AMMG.

